# Prefrontal stimulation prior to motor sequence learning alters multivoxel patterns in the striatum and the hippocampus

**DOI:** 10.1101/2021.07.01.450671

**Authors:** Mareike A. Gann, Bradley R. King, Nina Dolfen, Menno P. Veldman, Marco Davare, Stephan P. Swinnen, Dante Mantini, Edwin M. Robertson, Geneviève Albouy

**Affiliations:** Department of Movement Sciences, Movement Control and Neuroplasticity Research Group, KU Leuven, 3001 Leuven, Belgium; LBI - KU Leuven Brain Institute, KU Leuven, 3001 Leuven, Belgium; Department of Health and Kinesiology, College of Health, University of Utah, Salt Lake City, UT 84112, USA; Department of Clinical Sciences, College of Health and Life Sciences, Brunel University London, UB8 3PN Uxbridge, United Kingdom; Brain Imaging and Neural Dynamics Research Group, IRCCS San Camillo Hospital, 30126 Venice, Italy; Institute of Neuroscience and Psychology, University of Glasgow, G12 8QB Glasgow, United Kingdom

## Abstract

Motor sequence learning (MSL) is supported by dynamical interactions between hippocampal and striatal networks that are thought to be orchestrated by the prefrontal cortex. In the present study, we tested whether individually-tailored theta-burst stimulation of the dorsolateral prefrontal cortex (DLPFC) prior to MSL, can modulate multivoxel response patterns in the stimulated cortical area, the hippocampus and the striatum. Response patterns were assessed with multivoxel correlation structure analyses of functional magnetic resonance imaging data acquired during task practice and during resting-state scans before and after learning/stimulation. Results revealed that, across stimulation conditions, MSL induced greater modulation of task-related DLPFC multivoxel patterns than random practice. A similar learning-related modulatory effect was observed on sensorimotor putamen patterns under inhibitory stimulation. Furthermore, MSL as well as inhibitory stimulation affected (posterior) hippocampal multivoxel patterns at post-intervention rest. Exploratory analyses showed that MSL-related brain patterns in the posterior hippocampus persisted into post-learning rest preferentially after inhibitory stimulation. These results collectively show that prefrontal stimulation can alter multivoxel brain patterns in deep brain regions that are critical for the MSL process. They also suggest that stimulation influenced early offline consolidation processes as evidenced by a stimulation-induced modulation of the reinstatement of task pattern into post-learning wakeful rest.

## Introduction

The acquisition of new motor skills has been extensively studied using motor sequence learning (MSL) tasks during which participants integrate a series of movements into a temporally coherent structure^1,2^. This process is known to be supported by cortico-cerebellar, -striatal and -hippocampal networks^3–5^. Over the last decade, cortico-striatal and cortico-hippocampal networks have received particular attention and research has shown that the interaction between these systems - thought to be mediated by the dorsolateral prefrontal cortex (DLPFC) - is critical for the learning and retention of motor sequences^4,6,7^. As such, interventions designed to modulate hippocampal and striatal neural patterns are of great interest in the field of motor learning.

Inspired by the recent surge of research showing that non-invasive brain stimulation can be used as a tool to modulate brain responses in deeper regions (e.g.^8–11^), our group recently used repetitive transcranial magnetic stimulation (TMS) of the DLPFC in an effort to influence responses in deeper regions associated to motor learning^12^. Results indicated that inhibitory - as compared to facilitatory - DLPFC stimulation induced sustained connectivity in associative brain areas including the associative territories of the putamen and hippocampo-frontal networks during initial MSL. In contrast, connectivity patterns in the sensorimotor portions of the putamen appeared to be disrupted by inhibitory stimulation. Intriguingly, while frontal stimulation significantly modulated functional *connectivity* profiles, no effect of stimulation was observed on the level of *brain activity* of the striatum, the hippocampus or the DLPFC. This might suggest that prefrontal stimulation can modulate the connectivity of the target and deeper brain regions without inducing changes in activity levels. Alternatively, the examination of modulation of BOLD signal amplitude using univariate analyses might not be sensitive enough to highlight the subtle effect of stimulation on local brain patterns. Previous studies indeed suggest that univariate analyses might underestimate or even miss effects as they neglect potential relationships of activity profiles between voxels^13,14^. An avenue to address this issue is to employ multivariate analyses that are able to highlight activity patterns from multiple voxels and afford inferences about local brain patterns.

Research using multivariate approaches have effectively provided additional insights into the neural processes supporting MSL (e.g.^13–24^) as well as into TMS-related modulation of brain responses (e.g. ^8,19,25–31^). For example, Rafiei et al. (2021) were able to decode the applied TMS conditions by examining local brain patterns underneath the TMS coil while univariate analyses showed no stimulation-related activity changes in the same area. The vast majority of these multivariate investigations has employed representational analyses investigating whether brain patterns can discriminate e.g. different motor sequences or learning stages. An additional multivariate approach, that is particularly relevant in the context of learning and memory, is pairwise multivoxel correlation structure (MVCS^32,33^) analyses. Such an approach affords the opportunity to study the *similarity* of multivoxel brain patterns between different fMRI runs or sessions (e.g., at rest and/or during task practice). Using MVCS analyses, earlier studies have shown that local brain response patterns that were related to the learning episode can persist into subsequent rest periods (i.e., after learning has ended^32,34,35^). Importantly, the persistence of brain patterns is thought to reflect “replay” of learning-related neural patterns and to be functionally relevant as it can predict subsequent memory retention^32,34^.

In the present study, we used MVCS analyses to investigate whether DLPFC TMS can modulate local multivoxel brain patterns in the stimulated region as well as in the striatum and the hippocampus during MSL and at rest. Participants received either facilitatory or inhibitory theta-burst stimulation (i.e., intermittent iTBS and continuous cTBS, respectively^36^) of an individualized DLPFC target ***before*** performing a sequential serial reaction time task (SRTT^37^) or a control task (random SRTT; see Fig. 1). BOLD signal was acquired during task performance and during two resting-state runs administered pre-stimulation and post-task. We examined the effect of task (sequence vs. random) and stimulation (inhibitory vs. facilitatory) conditions on the task- and resting-state-related multivoxel patterns in four regions of interest (ROIs: DLPFC, hippocampus, associative putamen and sensorimotor putamen).

**Figure 1.**
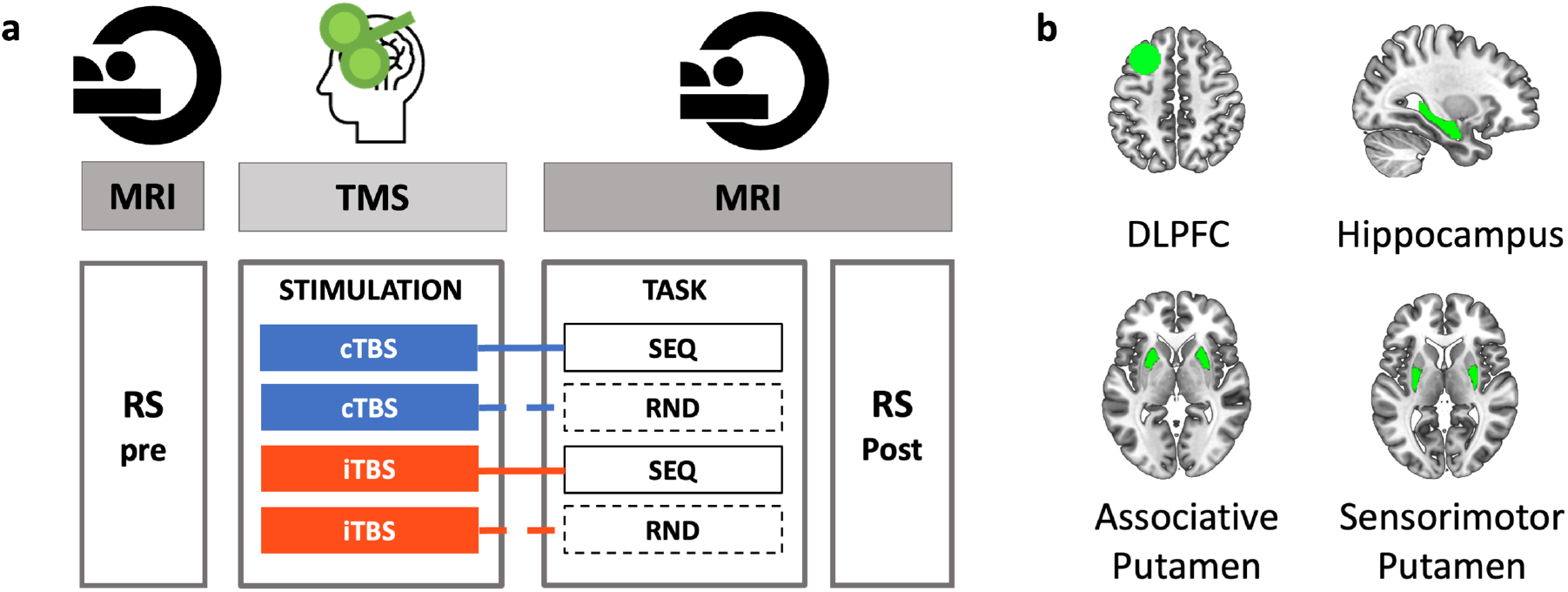
Experimental design and regions of interest. (a) In each experimental session, participants first underwent pre-TMS whole-brain resting-state (RS) fMRI scans, followed by T1-neuronavigated intermittent or continuous theta-burst stimulation (iTBS [190s] or cTBS [40s]) applied to an individually-defined DLPFC target outside the scanner (mean time between start RS pre and start TBS: 54.49min, range 43-77min). Directly following stimulation, participants were placed in the MR scanner where they were trained on the motor task (sequential [SEQ, indicated by solid lines] or random [RND, indicated by dashed lines] versions of the serial reaction time task) while BOLD images were acquired (mean delay between start TBS and start task: 15.71min, range 12–22min; mean duration of the task training: 11.5min, range 9.33–13.43min). After task completion, post-stimulation/task RS data were acquired (mean time between start TBS and start RS post: 32.49min, range 28–37min). The order of the four experimental conditions in this within-subject design [cTBS/SEQ (cSEQ), cTBS/RND (cRND), iTBS/SEQ (iSEQ), iTBS/RND (iRND)] was counterbalanced across participants. (b) Regions of interest (ROIs) for multivoxel correlation structure analyses. The dorsolateral prefrontal cortex (DLPFC) ROI consisted of a 10-mm-radius sphere centered around the individual TMS targets. The deeper ROIs were delineated using FSL FIRST and the Oxford-GSK-Imanova Striatal Connectivity Atlas. TMS – transcranial magnetic stimulation.

We expected the MVCS approach used in the current study to allow the detection of both learning- and stimulation-related modulation of brain responses in the ROIs *during task practice*. Specifically, we hypothesized that, across stimulation conditions, sequence learning would affect task-related multivoxel patterns more than random practice in all ROIs. As our earlier work suggests that inhibitory stimulation preferentially modulated functional responses in deep brain regions during MSL^12^, we predicted that inhibitory, as compared to facilitatory, stimulation would specifically affect these learning-related brain patterns. Additionally, based on the critical involvement of the hippocampus in motor sequence learning^4^ and on previous reports showing persistence of hippocampal patterns into rest after both declarative^8,32,34,38–40^ and motor^35^ learning, we predicted that motor sequence learning would induce greater hippocampal pattern persistence as compared to random practice. As inhibitory stimulation was expected to induce greater modulation of task-related patterns than facilitatory stimulation, we predicted that persistence of task patterns into post-learning rest would be stronger under inhibitory prefrontal stimulation.

## Material and Methods

The research presented in this manuscript is part of a larger experimental protocol (see Supplemental Fig. S1 for the full design). A subset of corresponding results is reported in ^12^.

### Ethics statement

Ethical approval was received from the local Ethics committee (UZ/KU Leuven, B322201525025). Written informed consent was provided by all participants before they participated in the study. Participants received compensation for their participation. The approved guidelines were respected for all procedures.

### Participants

Twenty-one young (range: 19-26years) right-handed^41^ participants took part in this study. Participants did not smoke or take psychoactive medications. They did not suffer from any known psychological, psychiatric (including anxiety^42^ and depression^43^) or neurological disorders and reported normal sleep during the month preceding the experiment (Pittsburgh Sleep Quality Index^44^). All participants had normal or corrected-to-normal vision, did not present any contra-indication for MRI or TMS, and were not considered musicians or professional typists. Two participants were excluded because of incidental findings on the acquired imaging data. The final sample included 19 participants (12 females; mean age=22.42±2.36). Yet, one participant did not complete one MR session (out of four) because of technical problems. Due to failure to appropriately perform the motor task (i.e., >3SD below the mean for accuracy), the behavioral and MRI data of two experimental sessions were excluded for another participant. One task fMRI run of a different participant and one post-task RS run of another participant were excluded due to excessive head motion (i.e., task: movement >2 voxels; RS: <100 remaining volumes after removal of volumes due to framewise displacement as described below). Depending on the conditions and contrasts tested, analyses therefore included 15 to 17 participants.

### General experimental procedure

The experimental protocol consisted of five sessions (one baseline and four TBS sessions). The TBS sessions took place at approximately the same time of day (±2h) and were separated by at least 6 days to avoid carry-over effects of the stimulation. Participants were instructed to refrain from alcohol consumption the day before and the day of the TBS sessions and to have a good night of sleep preceding each session (sleep quality and quantity assessed with the St. Mary’s Hospital Sleep Questionnaire^45^). Subjective vigilance at the time of testing was also assessed using the Stanford Sleepiness Scale^46^.

During the baseline session, participants were trained on a random version of the serial reaction time task (see below) to familiarize themselves with the task and equipment. A T1-weighted image as well as a RS functional run were acquired during this session. The four subsequent TBS sessions were organized according to a stimulation (2 levels: intermittent TBS [iTBS] vs. continuous TBS [cTBS]) by task (2 levels: sequence [SEQ] vs. random [RND]) within-subject design (Fig. 1a). In each TBS session, pre-TBS RS scans were first acquired before participants received T1-neuronavigated TBS on an individually-defined DLPFC target outside the scanner. Directly following stimulation, participants were placed back in the scanner where they were trained on the serial reaction time task. Post-stimulation/task RS scans were acquired immediately after task completion. The order of the four experimental conditions [cTBS/SEQ (cSEQ), cTBS/RND (cRND), iTBS/SEQ (iSEQ), iTBS/RND (iRND)] was counterbalanced across participants. However, due to participant exclusion (see details in the participants section), the different conditions were not perfectly balanced across visits in the analyzed data (see Supplemental Table S1). A factor including TBS visit order was therefore included in the behavioral and brain statistical analyses in order to control for this potential confounding factor. Note that the full experimental design included additional runs that are not reported in this manuscript (see Supplemental Fig. S1 for the full design).

### Serial reaction time task

Participants were trained on an explicit bimanual version of the serial reaction time task (SRTT^37,47^) implemented in the Psychophysics Toolbox in Matlab^48^ and routinely used in our group^49,50^. The task was performed on a specialized MR-compatible keyboard that was placed on the participant’s lap inside the MRI scanner. During the task, eight squares, representing the eight keys of the keyboard, were displayed on a screen that participants could see via a mirror when lying in the scanner. The color of the outline of the squares alternated between red and green, indicating rest and practice blocks, respectively. Each rest block lasted 15s during which participants were instructed to fixate on the screen without moving the fingers. At the start of the practice block, the outline of the squares turned to green and one of the eight squares filled green until the participants’ response was recorded. Participants were instructed to press the key corresponding with the green filled square with the corresponding finger (all fingers except thumbs) as quickly and as accurately as possible. As soon as the key was pressed, a new square changed to green, regardless of whether the response was correct or not (response to stimulus interval=0ms). Each practice block consisted of 48 key presses and each task session included 16 blocks of practice. The order in which the squares were colored green (and thus the order of finger movements) followed either a pseudorandom (RND) or a fixed, repeating sequential pattern (SEQ) according to the specific experimental condition. For the SEQ task condition, participants learned an eight-element sequence that was repeated six times per block. Two different sequences (4-7-3-8-6-2-5-1 and 7-2-8-4-1-6-3-5 with 1 representing the left little finger and 8 representing the right little finger, respectively) were used for the 2 different SEQ sessions. Note that due to experimental error, one participant learned the sequences 4-7-3-8-6-2-5-1 and 2-6-1-5-8-3-7-4 and another participant was trained on sequences 7-2-8-4-1-6-3-5 and 2-6-1-5-8-3-7-4. There was no repeating sequence in the RND task condition, but each square was filled once every eight elements (without repeating elements); consequently, each key was also pressed six times per block. Participants were informed whether the stimuli would follow a random or a repeating sequential pattern before the task started.

For each practice block, performance speed and accuracy were computed as the mean response time for correct responses (ms) and the percentage of correct responses, respectively. The behavioral data were analyzed with linear mixed models fitted in SPSS (IBM SPSS Statistics, Version 27, Armonk, NY, USA) using the performance speed or accuracy measures described above as the dependent variable. The TBS visit order (session 1-4), stimulation (2: cTBS/iTBS), block (16) and task (2: SEQ/RND) factors as well as the stimulation by task, stimulation by block, task by block and stimulation by task by block interactions were modelled as fixed effects. The experimental conditions (4) were modelled as repeated measures with the repeated covariance type ‘unstructured’ and the blocks (16) as repeated measures with the repeated covariance type ‘compound symmetry’ and both were combined in a Kronecker product covariance structure. Satterthwaite approximation for degrees of freedom adjustment was applied.

### TMS administration

#### Theta-burst stimulation

Theta-burst stimulation (TBS; a burst of 3 pulses given at 50Hz, repeated every 200ms^36^) was administered, outside the MRI scanner, on an individualized DLPFC target using a DuoMAG XT-100 rTMS stimulator (DEYMED Diagnostics s.r.o., Hronov, Czech Republic). TMS target identification was performed using a functionally-driven approach tailored to each individual^12^. Briefly, we first analyzed fMRI data from a sample of young healthy individuals^51^ independent from the sample of the current study and identified a cortical cluster functionally connected to both the striatum and hippocampus at rest^12^. The individual TMS targets in the current study were determined for each individual using baseline resting-state (RS) data as the coordinate showing maximal connectivity with both the hippocampus and striatum in a 15mm sphere centered around the DLPFC cluster identified above (−30 22 48mm^12^). The list of individual coordinates is reported in Supplemental Table S2. During DLPFC stimulation, the 70mm DuoMAG butterfly coil position was monitored online using neuronavigation (BrainSight, Rogue Research Inc, Montreal, Quebec, CA) and was placed at a 45° angle with the handle pointing posteriorly. Intermittent TBS (iTBS, 2s TBS trains repeated every 10s for 190s, 600 pulses) and continuous TBS (cTBS, 40s uninterrupted train of TBS, 600 pulses) were applied at 80% active motor threshold (aMT^36^). Motor evoked potentials measured with a belly-tendon EMG montage on the right flexor dorsal interosseous (FDI) muscle were used to (i) define the aMT, which was characterized during voluntary submaximal FDI contraction as the lowest intensity for which minimum 5/10 MEPs were distinguishable from background EMG^52,53^ and to (ii) assess cortico-spinal excitability pre- and post-stimulation (data not reported in the present manuscript but see Supplemental Fig. S1 for the full design).

### fMRI data acquisition and analysis

#### Acquisition

MR images were acquired on a Philips Achieva 3.0T MRI system equipped with a 32-channel head coil. During the baseline session, a high-resolution T1-weighted structural image was acquired using a MPRAGE sequence (TR/TE=9.6/4.6ms; voxel size=0.98×0.98×1.2mm^3^; FoV=250×250×228mm^3^; 190 coronal slices) for each participant.

RS fMRI data were acquired during the baseline session as well as during the four experimental sessions pre-TBS and post-task with an ascending gradient EPI pulse sequence for T2*-weighted images (TR/TE=1000/33ms; multiband factor 3; flip angle=80°; 42 transverse slices; interslice gap=.5mm; voxel size=2.15×2.14×3mm^3^; FoV=240×240×146.5mm^3^; matrix=112×110; 300 dynamic scans). A black screen (i.e., no visual stimuli) was shown during data acquisition. During the RS scans, participants were instructed to not move, to keep their eyes closed and to not think of anything in particular.

Task-related fMRI data were acquired in each experimental session with an ascending gradient EPI pulse sequence for T2*-weighted images (TR/TE=2000/29.8ms; multiband factor 2; flip angle=90°; 54 transverse slices; interslice gap=.2mm; voxel size=2.5×2.5×2.5mm^3^; FoV=210×210×145.6mm^3^; matrix=84×82; 345.09±22.37 dynamical scans).

#### Analyses

##### Pre-processing

The fMRI data were preprocessed using SPM12 (http://www.fil.ion.ucl.ac.uk/spm/software/spm12/; Wellcome Centre for Human Neuroimaging, London, UK). The reoriented structural image was segmented into gray matter (GM), white matter (WM), cerebrospinal fluid (CSF), bone, soft tissue, and background. Task-based and RS functional volumes of each participant were first slice-time corrected (reference: middle slice). Images were then realigned to the first image of each session and, in a second step, realigned to the mean functional image computed across all the individuals’ fMRI runs using rigid body transformations. The mean functional image was co-registered to the high-resolution T1-weighted anatomical image using a rigid body transformation optimized to maximize the normalized mutual information between the two images. The resulting coregistration parameters were then applied to the realigned functional images. To optimize voxel pattern analyses, functional and anatomical data remained in subject-specific (i.e., native) space, and no spatial smoothing was applied to functional images^32,35^.

##### ROI definition

Four ROIs were considered in the main analyses (see Fig. 1b) and consisted of the individual left DLPFC target, the bilateral hippocampus, as well as the bilateral associative and sensorimotor parts of the putamen. The left DLPFC ROI was defined at the individual level using Marsbar (http://marsbar.sourceforge.net^54^) as a 10mm-radius sphere centered around the individual TMS target (see Supplemental Table S2 for the list of coordinates). Note that as the individual DLPFC coordinates were provided in MNI space^12^, the DLPFC ROIs were mapped back to native space using the individual’s inverse deformation field output from the segmentation of the anatomical image. The hippocampus and putamen ROI were created in the native space of each individual using the FMRIB’s Integrated Registration Segmentation Toolkit (FSL FIRST; http://fsl.fmrib.ox.ac.uk/fsl/fslwiki/FIRST). Based on the well-described functional organization of the putamen in sub-territories^55–57^ and on previous evidence showing that the effect of DLPFC TBS on putamen functional responses is different between these sub-regions^12^, the putamen ROI was split into associative and sensorimotor territories in the present study. To do so, masks of the sensorimotor and associative (executive) portions of the putamen were created using the Oxford-GSK-Imanova Striatal Connectivity Atlas (striatum-con-label-thr25/50-3sub; https://fsl.fmrib.ox.ac.uk/fsl/fslwiki/Atlases/striatumconn; FSL). These MNI-space-based masks were reverse transformed into each participant’s native space and applied to the individual’s putamen segment output from FSL FIRST. Additionally, we performed exploratory analyses on anterior and posterior hippocampal sub-territories obtained after a split of the individual hippocampal segment at the middle slice of the coronal extension. These analyses are considered exploratory as it remains unknown whether the well-described antero-posterior functional organization of the hippocampus^58,59^ applies to motor functions.

##### Multi-voxel correlation structure (MVCS) analyses

The analysis pipeline was written in Matlab and followed similar procedures as in^35^. Prior to running MVCS analyses, additional preprocessing of the time series was completed. Specifically, whole-brain signal was detrended and high-pass filtered (cutoff=1/128). For the analyses including runs acquired with different TRs (i.e., analyses including both RS and task runs), runs with the faster TR were down sampled to the lower TR (here RS data were down sampled to the task TR). Furthermore, if the framewise displacement of any given volume exceeded 0.5mm, that volume as well as the subsequent one were excluded (5.35% of volumes were excluded for task runs, 1.27% for RS runs and 2.86% for analyses combining task and RS runs). At the ROI level, only voxels with >10% GM probability were included in the analyses. Regression analyses were performed on the fMRI time-series of the remaining voxels in each ROIs to remove nuisance factors. The three first principal components of the signal extracted from the WM and CSF masks created during segmentation of the anatomical image were regressed out from the time series (6 regressors in total). The 6-dimensional head motion realignment parameters, as well as the realignment parameters squared, their derivatives, and the squared derivatives, were also used as regressors (24 regressors in total). Lastly, the number of volumes was matched across runs included in the specific pair of runs included in that particular analysis. To do so, in the run of the analyzed pair presenting a higher number of scans, we selected the x volumes around the middle volume of the run, with x defined as the number of volumes in the run with the smaller number of scans.

Multi-voxel correlation structure (MVCS) matrices were computed for each ROI and each run with similar procedures as in previous research^32,34,35,60^. Specifically, Pearson’s correlations were computed between each of n BOLD-fMRI voxel time courses, yielding an n by n MVCS matrix per ROI and per run. Pearson’s correlation coefficients were then Fisher Z-transformed to ensure normality. A similarity index (SI) reflecting the similarity of the multivoxel patterns between two specific fMRI runs (see Fig. 2 for an example) was computed as the r-to-z transformed correlation between the two MVCS matrices of interest^32^. SI were computed between (1) early and late task practice (i.e., first and second half of task practice) in order to assess the effect of task practice on brain patterns, (2) RS pre and RS post in order to assess the effect of the experimental interventions on brain patterns at rest, and (3) task practice and RS post in order to investigate whether task-related brain patterns persisted during subsequent rest. Importantly, in order to test whether pattern similarity was influenced by our experimental interventions (i.e. task and stimulation conditions), SI were compared, for each ROI, using linear mixed modelling. Models were fitted in SPSS (IBM SPSS Statistics, Version 27, Armonk, NY, USA) with the TBS visit order (1-4), stimulation (2: cTBS/iTBS) and task (2: SEQ/RND) factors as well as with the stimulation by task interaction as fixed effects. The experimental conditions (4) were indicated as repeated measures with the repeated covariance type ‘unstructured’ and we used Satterthwaite approximation for degrees of freedom adjustment. SI means (adjusted and unadjusted for the TBS visit fixed effect) can be found for all ROIs in Supplemental Table S3. Results of these analyses were considered significant if *p*<.05. Results surviving false-discovery-rate (FDR^61^) correction for multiple comparisons are indicated with an asterisk in the corresponding tables.

**Figure 2.**
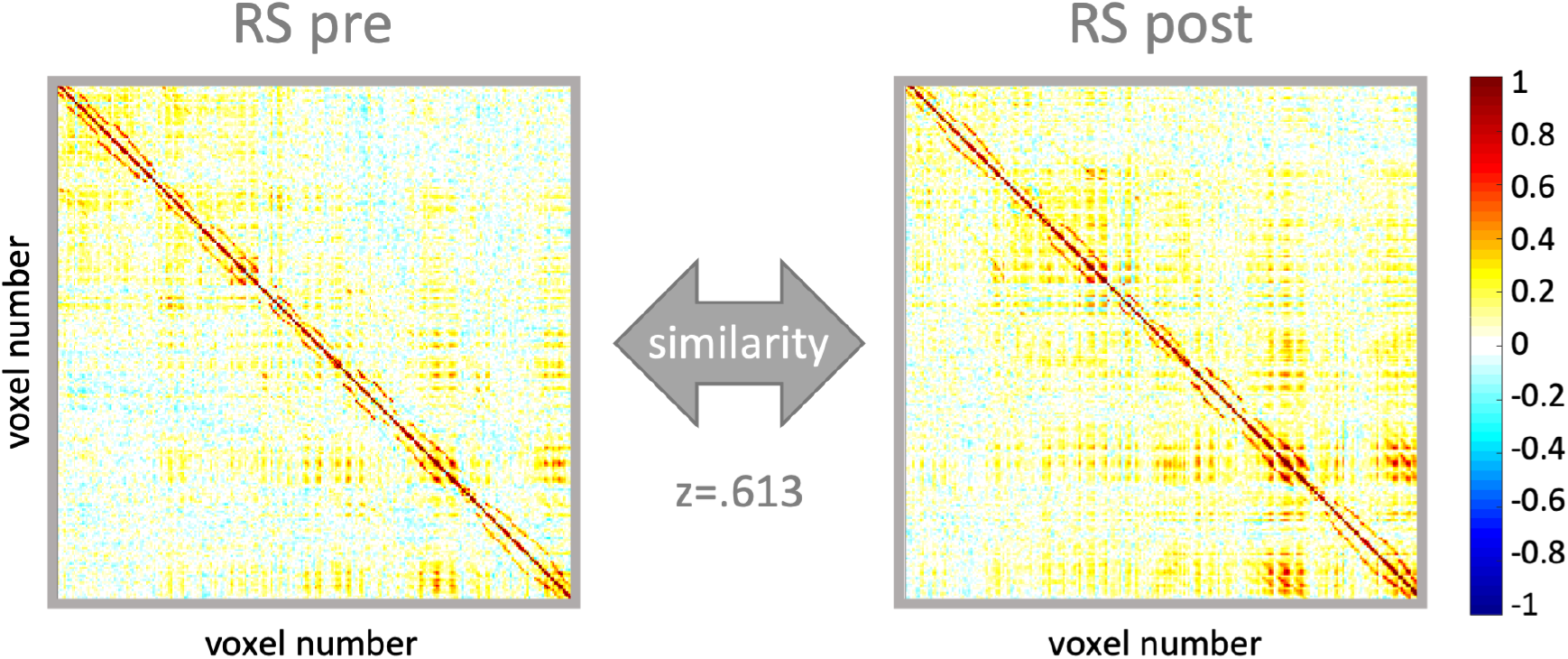
Multivoxel correlation structure (MVCS) for an exemplar ROI and participant. Each matrix depicts the correlation between each of the n voxels of the ROI with all the other voxels of the ROI during a specific fMRI run. The similarity between two matrices (here pre and post resting-state (RS) runs) is calculated as the r-to-z transformed correlation between the two MVCS. Resulting Z scores are compared between task and stimulation conditions.

## Results

### Behavior

Performance speed and accuracy (see Fig. 3) were analyzed using linear mixed models including task (2: SEQ/RND), stimulation (2: cTBS/iTBS), block (16) and TBS visit order (session 1-4) as factors. Analyses of performance speed showed a main effect of task (F_(1,49.558)_=40.794, *p*<.001) whereby performance on the SEQ condition was better than on the RND condition. A main effect of block was observed (F_(15,264.308)_=17.659, *p*<.001) indicating that participants became faster as a function of practice. The block-to-block speed improvement was stronger in the SEQ than in the RND task (task by block interaction; F_(15,282.182)_=24.057, *p*<.001). The main effect of stimulation (F_(1,18.67)_=.587, *p*=.447) as well as the stimulation by block (F_(15,279.893)_=.522, *p*=.928) and stimulation by task by block interactions (F_(15,274.729)_=.657, *p*=.826) were not significant, but a trend for a stimulation by task interaction was observed (F_(1,49.522)_=3.629, *p*=.063).

**Figure 3.**
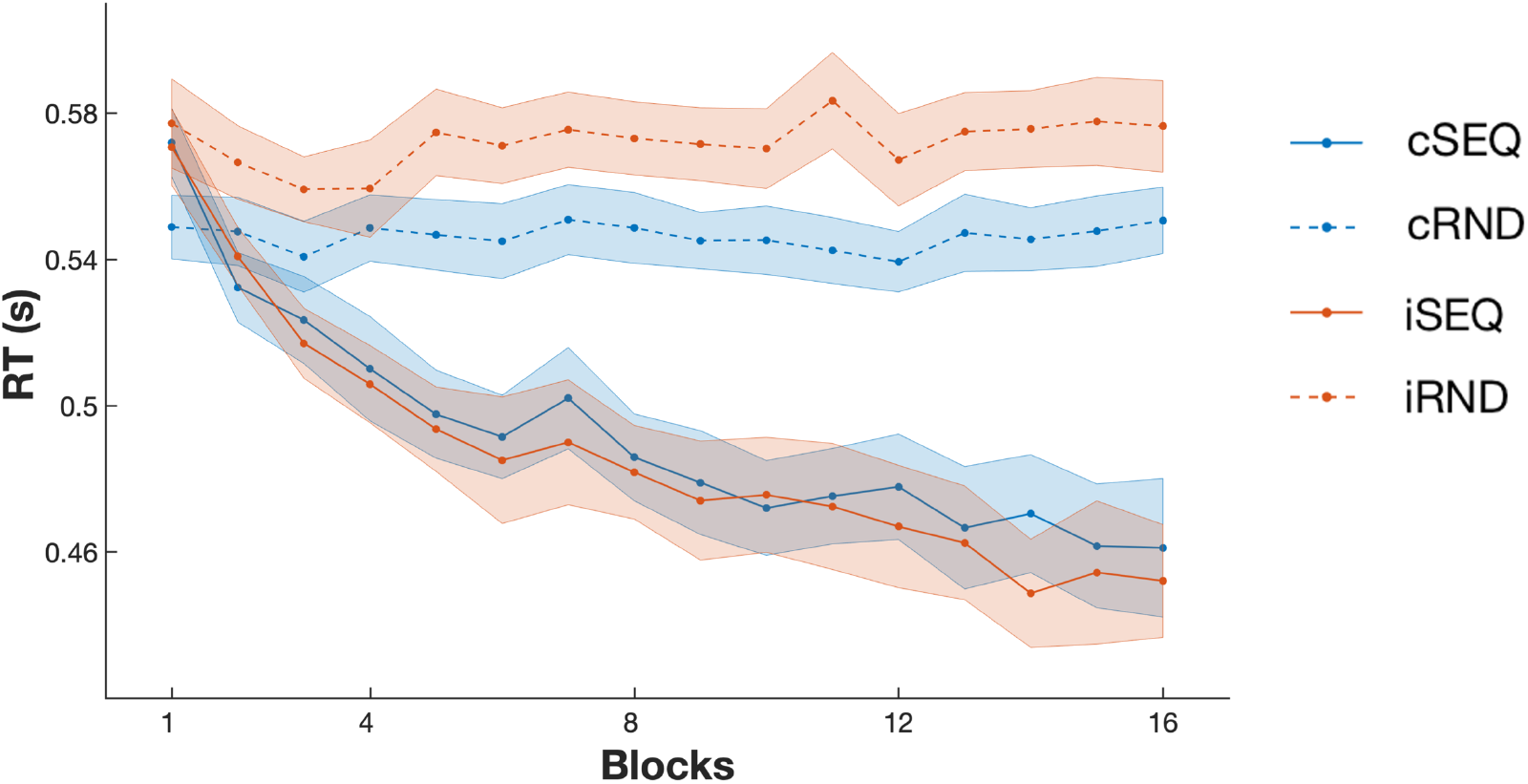
Behavioral results. Performance speed (reaction time, RT) improved over the course of training in the sequence task (SEQ) conditions and stayed stable in random task (RND) conditions. The stimulation intervention (c: continuous and i: intermittent) did not affect motor performance nor motor learning. Lines and dots represent average RT per block and shaded areas represent SEM.

Analyses of performance accuracy revealed a trend for a task effect (F_(1,41.118)_=3.63, *p*=.064) as well as for a block effect (F_(15,201.325)_=1.557, *p*=.088) but no significant task by block interaction (F_(15,243.436)_=.868, *p*=.6), main effect of stimulation (F_(1,40.437)_=1.251, *p*=.27), stimulation by task interaction (F_(1,39.176)_=.318, *p*=.576), stimulation by block interaction (F_(15,235.442)_=.849, *p*=.622) nor stimulation by task by block effect (F_(15,241.086)_=1.09, *p*=.367) were observed.

Altogether, the behavioral results showed that participants learned the motor sequence and that the stimulation intervention did not significantly influence motor sequence learning nor overall motor performance.

### MVCS

#### Pattern similarity between early and late task practice stages

Linear mixed models tested whether task and stimulation conditions influenced the similarity of task-related multivoxel patterns between early and late stages of practice for each ROI (see Table 1 for statistics).

**Table 1.** Task by stimulation effects on pattern similarity between early and late task practice stages controlling for the visit effect.

|  | dfs | F | p |
| --- | --- | --- | --- |
| **DLPFC** |  |  |  |
| *Task effect* | 1,18.141 | 5.137 | **.036** |
| *Stimulation effect* | 1,15.682 | 1.717 | .209 |
| *Interaction* | 1,14.791 | 4.031 | .063 |
| **Hippocampus** |  |  |  |
| *Task effect* | 1,14.662 | 3.087 | .100 |
| *Stimulation effect* | 1,15.381 | .162 | .692 |
| *Interaction* | 1,14.713 | .201 | .660 |
| **Associative Putamen** |  |  |  |
| *Task effect* | 1,17.443 | .315 | .582 |
| *Stimulation effect* | 1,16.970 | 1.630 | .219 |
| *Interaction* | 1,13.405 | 2.366 | .147 |
| **Sensorimotor Putamen** |  |  |  |
| *Task effect* | 1,15.288 | .245 | .628 |
| *Stimulation effect* | 1,18.201 | <.001 | .992 |
| *Interaction* | 1,8.403 | 10.990 | **.01\*** |

Bold values indicate $p<.05$ . False-discovery-rate (FDR) correction for multiple comparisons (4 ROIs) was applied with the Benjamini-Hochberg procedure<sup>61</sup>. Asterisk (\*) indicates significance at $p_{FDR}<.05$ . DLPFC – dorsolateral prefrontal cortex, dfs – degrees of freedom.

Results show that DLPFC pattern similarity between early and late stages of task practice were influenced by the task condition (F_(1,18.141)_=5.137, *p_unc_*=.036; Fig. 4a). Specifically, DLPFC patterns were less similar between the early and late task stages in the sequence as compared to the random task conditions. No stimulation effect and a trend for a task by stimulation interaction were observed. Altogether, these results suggest that, across stimulation conditions, DLPFC multivoxel patterns were affected by motor sequence learning more than by random practice.

**Figure 4.**
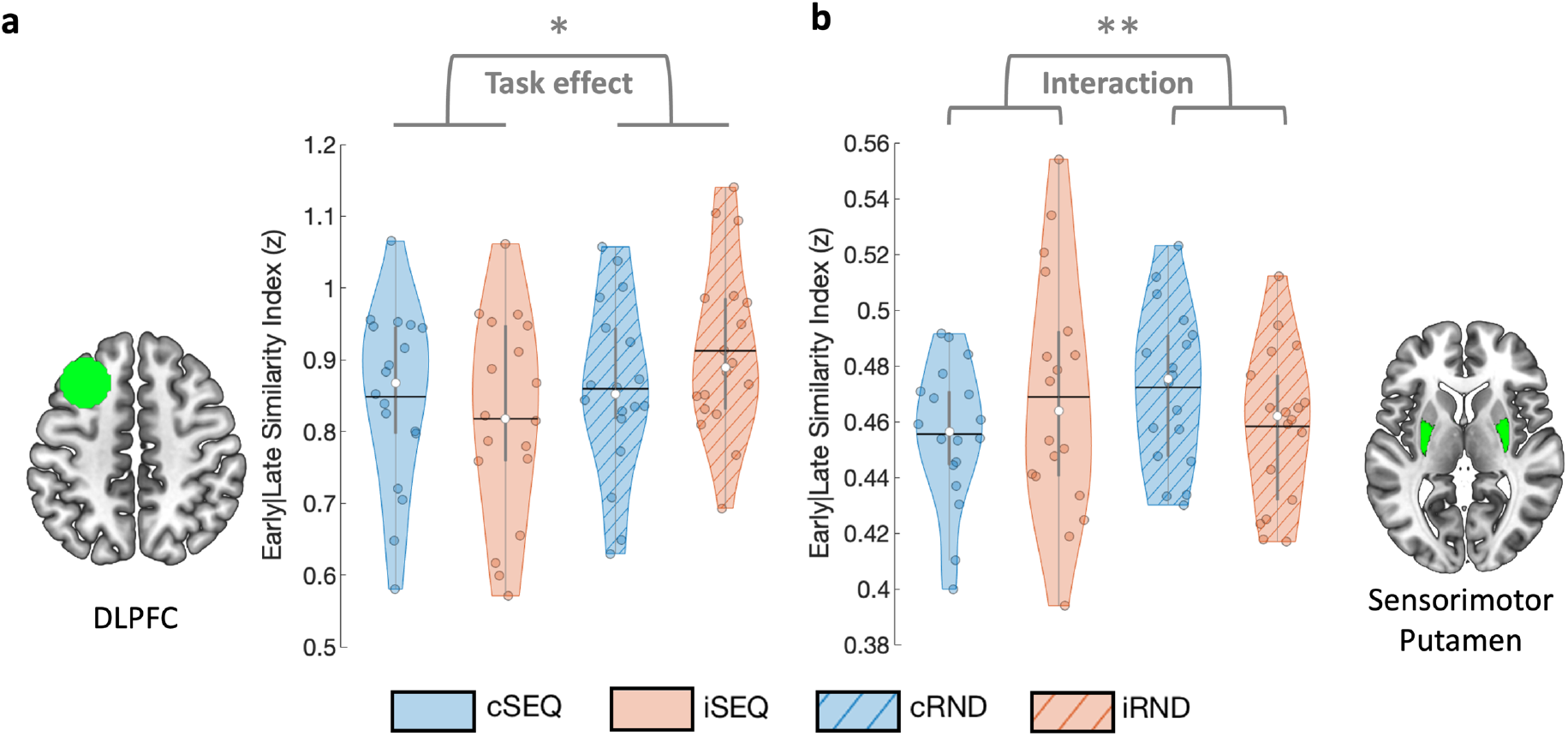
Similarity between early and late task practice stages. (a) DLPFC pattern similarity between early and late task practice was influenced by task condition such that sequence learning resulted in lower early-late similarity as compared to random practice. (b) The sensorimotor part of the putamen showed a significant task by stimulation interaction whereby early-late similarity was lower during sequence as compared to random practice, after cTBS as compared to iTBS. Colored circles represent individual data, jittered in arbitrary distances on the x-axis within the respective violin plot to increase perceptibility. Black horizontal lines represent means and white circles represent medians. The shape of the violin plots depicts the distribution of the data and grey vertical lines represent quartiles. Asterisk indicates significance at *p*<.05 (*) and at *pFDR*<.05 (**). Violin plots were created with^62^. DLPFC – dorsolateral prefrontal cortex, SEQ – sequence learning task version, RND – random task version, c – continuous, i – intermittent.

Sensorimotor putamen pattern similarity between early and late stages of task practice (see Fig. 4b) were not influenced by task or stimulation conditions. However, a significant task by stimulation interaction effect (F_(1,8.203)_=10.99, *p_unc_*=.01) was observed whereby early-late similarity was lower during sequence as compared to random practice, after cTBS as compared to iTBS. Specifically, follow-up analyses indicated that early-late similarity was significantly lower in cSEQ as compared to cRND conditions (F_(1,7.588)_=11.713, *p_unc_*=.01). These results suggest that sequence learning specifically affected sensorimotor putamen patterns under the influence of inhibitory cTBS of the DLPFC.

No significant effects were observed in the associative putamen and the hippocampus (all uncorrected *p*s>.05, see Table 1).

#### Pattern similarity between RS pre- and post-intervention

Linear mixed models tested whether task and stimulation conditions influenced the similarity of multivoxel patterns during resting-state measured pre- and post-intervention (see Table 2). In the hippocampus, pattern similarity at rest between pre- and post-intervention runs (see Fig. 5) was influenced by the task condition (F_(1,18.429)_=6.949, *p_unc_*=.017). Specifically, across stimulation conditions, hippocampal patterns were less similar between the pre- and post-measurements after SEQ as compared to RND task practice. Additionally, hippocampal multivoxel patterns at rest were also influenced by the stimulation condition (F_(1,14.319)_=6.837, *p_unc_*=.019) whereby lower similarity between pre- and post-RS was observed in the cTBS as compared to the iTBS conditions. No task by stimulation interaction effect was observed. Exploratory analyses on sub-territories of the hippocampus showed that the effects described above were driven by the posterior portion of the hippocampus in which the stimulation as well as the task effects were also observed (stimulation effect: F_(1,11.878)_=7.62, *p_unc_*=.017; task effect: F_(1,18.879)_=6.239, *p_unc_*=.022; Supplemental Table S4, Supplemental Fig. S2). No such effects were observed in the anterior hippocampus. Collectively, these results suggest that motor sequence learning as well as inhibitory cTBS of the DLPFC affected (posterior) hippocampal multivoxel patterns measured during rest post-intervention.

**Table 2.** Task by stimulation effects on pattern similarity between RS pre- and post-stimulation/task controlling for the visit effect.

|  | dfs | F | *p* |
| --- | --- | --- | --- |
| **DLPFC** |  |  |  |
| *Task effect* | 1,18.068 | .016 | .900 |
| *Stimulation effect* | 1,16.770 | .061 | .808 |
| *Interaction* | 1,16.904 | 1.554 | .230 |
| **Hippocampus** |  |  |  |
| *Task effect* | 1,18.429 | 6.949 | **.017** |
| *Stimulation effect* | 1,14.319 | 6.837 | **.019** |
| *Interaction* | 1,17.480 | .773 | .391 |
| **Associative Putamen** |  |  |  |
| *Task effect* | 1,16.348 | 1.950 | .181 |
| *Stimulation effect* | 1,16.309 | 3.003 | .102 |
| *Interaction* | 1,14.025 | .065 | .803 |
| **Sensorimotor Putamen** |  |  |  |
| *Task effect* | 1,18.214 | 2.171 | .158 |
| *Stimulation effect* | 1,17.271 | .276 | .606 |
| *Interaction* | 1,16.990 | .054 | .819 |

*Bold values indicate $p < .05$ . None of the results presented in this table survived FDR correction for 4 ROIs. DLPFC – dorsolateral prefrontal cortex, dfs – degrees of freedom.*

**Figure 5.**
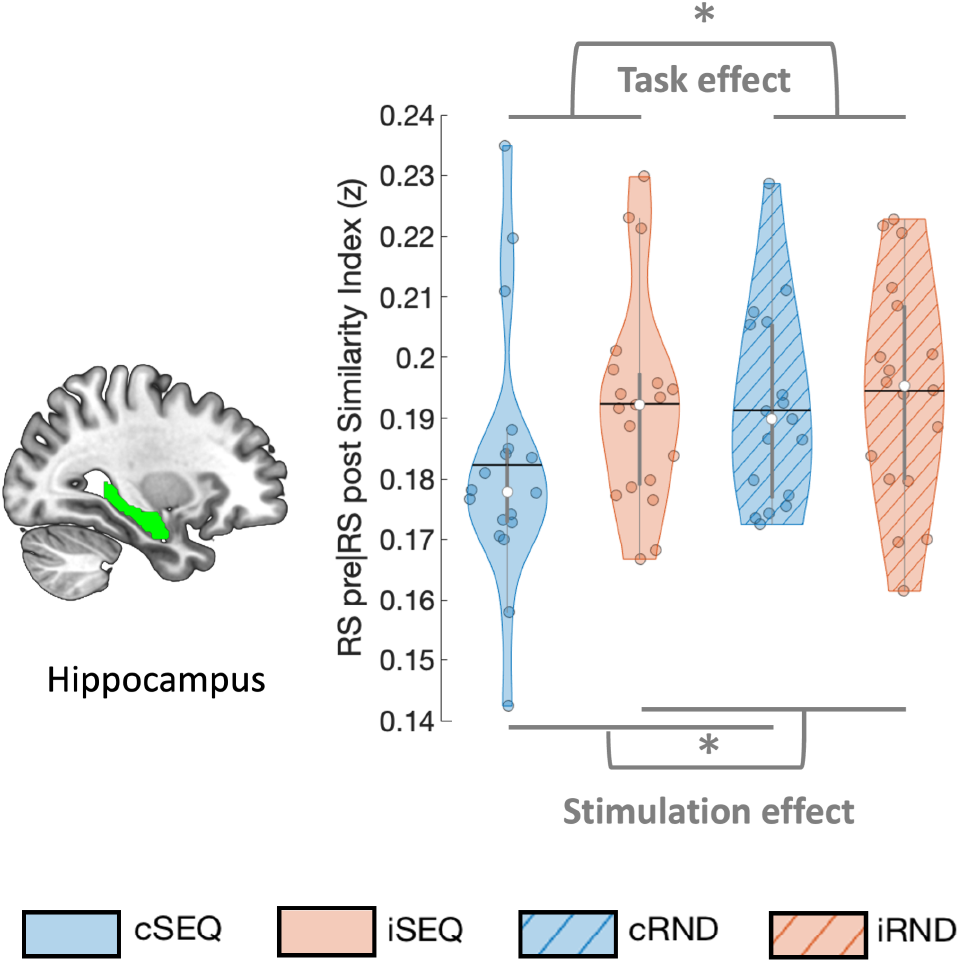
Similarity between RS pre- and post-intervention. Hippocampal pattern similarity between pre and post RS was influenced by task and stimulation condition such that lower similarity was observed after sequence learning as compared to random practice as well as after cTBS as compared to iTBS. Colored circles represent individual data, jittered in arbitrary distances on the x-axis within the respective violin plot to increase perceptibility. Black horizontal lines represent means and white circles represent medians. The shape of the violin plots depicts the distribution of the data and grey vertical lines represent quartiles. Asterisk (*) indicates significance at *p*<.05. RS – resting-state, SEQ – sequence learning task version, RND – random task version, c – continuous, i – intermittent.

No significant effects were observed on the resting-state similarity indexes of the DLPFC, associative or sensorimotor putamen (all uncorrected *p*s>.05, see Table 2).

#### Pattern similarity between task practice and RS post-intervention

Linear mixed models tested whether task and stimulation conditions influenced the persistence of multivoxel brain patterns from task into post-task RS runs (see Table 3).

**Table 3.** Task by stimulation effects on pattern similarity between task practice and RS post stimulation/task.

|  | dfs | F | p |
| --- | --- | --- | --- |
| **DLPFC** |  |  |  |
| *Task effect* | 1,18.914 | 2.177 | .157 |
| *Stimulation effect* | 1,15.338 | .084 | .776 |
| *Interaction* | 1,17.641 | 3.186 | .091 |
| **Hippocampus** |  |  |  |
| *Task effect* | 1,18.400 | .602 | .448 |
| *Stimulation effect* | 1,15.031 | .046 | .833 |
| *Interaction* | 1,18.068 | 1.267 | .275 |
| **Associative Putamen** |  |  |  |
| *Task effect* | 1,17.212 | .583 | .456 |
| *Stimulation effect* | 1,14.717 | 1.151 | .301 |
| *Interaction* | 1,14.879 | .517 | .483 |
| **Sensorimotor Putamen** |  |  |  |
| *Task effect* | 1,15.389 | .305 | .588 |
| *Stimulation effect* | 1,15.141 | 2.028 | .175 |
| *Interaction* | 1,18.379 | 1.945 | .180 |

*DLPFC – dorsolateral prefrontal cortex, dfs – degrees of freedom.*

Pattern similarity between these runs was not influenced by the task or stimulation conditions in any of the ROIs. However, exploratory analyses on hippocampal sub-territories showed a stimulation effect in the posterior hippocampus (F_(1,15.074)_=10.336, *p_unc_*=.006; Supplemental Table S5, Supplemental Fig. S2) with higher similarity in the cTBS as compared to the iTBS conditions. A stimulation by task interaction (F_(1,16.333)_=4.535, *p_unc_*=.049) was also observed whereby the difference in similarity between the stimulation conditions was larger in the sequence as compared to the random task. Specifically, follow-up analyses indicate that similarity between task practice and RS post was significantly higher in cSEQ as compared to iSEQ conditions (F_(1,15.173)_=9.280, *p_unc_*=.005). No task effect was observed.

No significant effects were observed in the anterior part of the hippocampus (all uncorrected *p*s>.05, see Supplemental Table S5).

## Discussion

In this study, we used multivariate correlation structure analyses to investigate the effects of prefrontal stimulation as well as motor sequence learning (MSL) on brain patterns in the DLPFC, hippocampus and putamen. While the behavioral data showed an expected task effect, the stimulation did not affect motor performance. At the brain level, we observed both task and stimulation-related modulation of multivoxel brain patterns. Specifically, our data indicate that initial motor sequence learning, irrespective of the stimulation condition, induced greater modulation of task-related DLPFC multivoxel patterns than random practice. A similar learning-related modulatory effect was observed on sensorimotor putamen patterns under inhibitory as compared to facilitatory stimulation. Furthermore, sequence learning as well as inhibitory DLPFC stimulation affected (posterior) hippocampal multivoxel patterns measured at rest after the intervention. Exploratory analyses suggest that these pattern modulations resulted from higher persistence of task-related posterior hippocampus pattern into subsequent rest after sequence learning as compared to random practice and under inhibitory as compared to facilitatory stimulation. Altogether, our findings indicate that motor sequence learning and inhibitory DLPFC stimulation can alter brain patterns in the stimulated target and in deeper regions during both task practice and at rest. They also suggest that stimulation influenced early motor memory consolidation during wakefulness as evidenced by stimulation-induced modulation of the reinstatement of task pattern into rest immediately following learning.

### Sequence learning modulated task-related patterns in the DLPFC

Brain imaging analyses revealed that initial motor sequence learning altered task-related DLPFC patterns more than random task practice irrespective of stimulation condition. Specifically, left DLPFC patterns were less similar between early and late task practice in the sequence than in the random condition. These results concur with previous reports of DLPFC activation during (early stages of) MSL^4,7,57,63–66^, and more particularly with evidence of preferential DLPFC recruitment during sequence learning as compared to random practice^65,66^. In the same vein, a recent study showed motor sequence representation in the DLPFC, such that multivariate patterns could effectively discriminate different motor sequences irrespective of the learning stage^24^. Interestingly, although the DLPFC has traditionally been associated with early MSL stages, multivariate approaches also highlighted the ability of the DLPFC to discriminate different *consolidated* motor sequences^24^ and *consolidated vs. newly learned* motor sequences^17^. Altogether, these studies suggest that DLPFC patterns carry sequence representation and that these representations are modulated by the learning stage. Our results are in line with these observations and suggest sequence-specific pattern modulation in the DLPFC during *initial* MSL.

Against our expectations, we observed no effect of stimulation on multivoxel patterns of the targeted DLPFC. These results are in conflict with previous studies showing TMS effects on the multivariate brain patterns of various cortical targets - including the DLPFC - during resting-state but also during task practice^8,25–27,30^. It remains unclear why no such effects were observed in the current study. Several methodological differences between our work and previous research might have contributed to these discrepancies (e.g., offline vs. online TMS^30^; group vs. individual analyses^26^; MVCS vs. classification^8,19,25^; control task vs. control stimulation^8,25,27^) but future research is certainly warranted to examine this question systematically.

In conclusion, the present multivariate approach highlighted sequence-specific modulation of DLPFC patterns that were not observed with our previous univariate analyses^12^. However, results did not reveal any stimulation effect on the multivoxel pattern of the TMS target.

### Sequence learning modulated task-related patterns in the sensorimotor putamen under inhibitory prefrontal stimulation

Our data showed that sequence learning affected task-related patterns in the bilateral sensorimotor putamen more than random practice and that this effect was more pronounced under inhibitory as compared to facilitatory stimulation. These results were mainly driven by a larger change in pattern between early and late stages of initial motor sequence learning under inhibitory stimulation. Interestingly, the effect of task condition - observed in the current study under the effect of inhibitory stimulation - is in line with previous research reporting sequence-learning-related modulation of striatal multivariate patterns^14,18,24^. Specifically, these recent studies have shown that (sensorimotor) striatal patterns can not only discriminate different motor sequences^18^ but also different motor sequence learning stages (e.g., new vs. learned sequences^14,24^; early versus late stages of initial learning^18^), although some of these effects might only be observed during full speed as compared to paced motor performance^14^. Our results confirm these previous findings and suggest that learning-related modulation of sensorimotor putamen patterns are preferentially observed under inhibitory stimulation condition. However, pattern similarity analyses do not allow us to conclude on whether stimulation *potentiated* or *disrupted* the learning-related modulation of sensorimotor putamen patterns. Based on our previous work suggesting that inhibitory stimulation *disrupted* the increase in sensorimotor putamen recruitment usually observed over the course of initial learning^12^, we speculate that inhibitory stimulation might have disrupted sensorimotor putamen patterns in the course of MSL. Although the multivoxel patterns in the bilateral associative putamen showed similar task by stimulation modulation as the sensorimotor part, these effects did not reach significance (see Supplemental Table S3). This stands in contrast with previous reports of motor sequence learning-related modulations of associative striatum multivariate patterns^14,18^ and with our earlier work suggesting that inhibitory stimulation resulted in greater maintenance of associative striatum recruitment during MSL^12^. It is worth noting, however, that most of the effects discussed above were observed in the caudate nucleus rather than in the associative portion of the putamen investigated in the current study. It would be worthwhile for future research to examine whether motor sequence learning differently affects pattern in these sub-regions of the associative striatum.

Altogether, the multivariate approach used in this study highlighted similar results as in our previous univariate work for the sensorimotor putamen but did not provide evidence of such modulatory effects for the associative putamen.

### Sequence learning and inhibitory prefrontal stimulation modulated hippocampal patterns during post-intervention rest

Our results indicate that sequence learning as well as inhibitory prefrontal stimulation modulated resting-state patterns in the bilateral (posterior) hippocampus. Specifically, pre- and post-intervention resting-state hippocampal patterns were more dissimilar after inhibitory (as compared to facilitatory) stimulation and after sequence learning (as compared to random task practice). In line with these findings, exploratory analyses on the posterior portion of the bilateral hippocampus revealed stronger persistence of task-related brain patterns into post-task rest after inhibitory stimulation as compared to facilitatory stimulation, and this more for the sequence condition as compared to the random condition. These findings suggest that the pre-post pattern dissimilarity described above might be the result of persistence of MSL-related brain patterns into rest and that such pattern persistence was specifically potentiated by inhibitory stimulation.

Persistence of task-related hippocampal patterns into post-learning wakeful rest immediately following learning has been consistently observed in the declarative memory domain^8,32–34,38–40^ and only more recently reported in the motor memory domain^35^. It is argued that such persistence reflects “reactivation” or “replay” of patterns that were previously expressed during learning^33,35^. Importantly, these reactivations are thought to be critical for memory processing as there is evidence that pattern persistence during post-learning rest is related to subsequent consolidation^32,34,38,39^. The present findings extend this previous research and provide critical evidence that hippocampal multivariate patterns related to initial motor sequence learning, but not to random task practice, persist into immediate post-learning rest. Interestingly, we observed pattern persistence in the posterior but not in the anterior portion of the hippocampus. These results are in line with traditional views, mainly based on rodent work, suggesting that the posterior hippocampus is implicated in cognitive functions such as learning, memory and spatial navigation while the anterior portion of the hippocampus is rather associated to stress and anxiety-related behaviors (^67^ but see e.g.^59,68^ for revised views on hippocampal organization). The current data extend these observations to motor learning and point towards a preferential role of the posterior hippocampus in the early motor memory consolidation process.

Importantly, this study demonstrates that these post-learning hippocampal pattern reinstatements are modulated by prefrontal stimulation. Specifically, the sequence-learning-related persistence effect was higher after inhibitory as compared to facilitatory DLPFC stimulation. This stimulation effect is in line with previous work from our group showing prolonged recruitment of fronto-(posterior) hippocampal networks after inhibitory stimulation during motor sequence learning^12^. We argue that these stimulation-induced modulations of hippocampo-frontal connectivity might have altered brain patterns during task practice, which would, in turn, influence patterns during post-learning rest. This interpretation remains however speculative as we did not observe modulations of hippocampal patterns during task practice in the current study.

Collectively, our results are the first to show that the persistence of (posterior) hippocampal multivariate patterns into subsequent rest was influenced by motor sequence learning as well as by prefrontal stimulation. Future research will investigate whether such task and stimulation-induced modulations of pattern persistence can influence the subsequent motor memory consolidation process as it was described in the declarative memory domain^32,34,38,39^.

### Prefrontal stimulation did not affect motor performance

In contrast to previous studies applying disruptive prefrontal stimulation prior or during MSL^69–72^ our results did not reveal any significant effects of stimulation on motor performance. These discrepancies might arise from several differences in methodology, such as the presence of reward during task practice, the awareness of the sequential material to learn (implicit / explicit learning), the task features (e.g., unimanual / bimanual) or the specific stimulation patterns used (1Hz, 5Hz rTMS and single pulse TMS / TBS). Interestingly, as the observed stimulation-specific modulations in multivoxel patterns of the hippocampus and putamen did not induce behavioral differences, it is tempting to speculate that such pattern modulations reflect compensatory brain mechanisms that maintained behavioral performance at similar levels between stimulation conditions. Specifically, one could have expected that the potential disruptive effect of inhibitory DLPFC stimulation observed during motor sequence learning on sensorimotor putamen patterns would result in poorer performance. As no differences in motor behavior were observed between stimulation conditions, we propose that the sustained engagement of (associative) hippocampo-frontal areas observed during sequence learning under inhibitory stimulation^12^ – and presumably reflected in the present study by higher hippocampal pattern persistence during post-learning rest – might have counteracted the negative effect of inhibitory stimulation on sensorimotor putamen patterns as well as on motor performance. This interpretation, however, remains hypothetical.

### Considerations

The current experiment did not include sham stimulation conditions. Due to ongoing debates in the literature with respect to the appropriateness of sham stimulation for within-subject TMS protocols^73,74^, we prioritized the inclusion of a control task condition rather than a control stimulation condition. The random task condition allowed us to test for sequence learning-specific effects and to disentangle whether stimulation effects depend on the task state under which stimulation was active (i.e., learning vs. control). However, the lack of a sham control condition induced limitations, as any significant stimulation-specific result in the current study is derived from comparisons between two active stimulation conditions and not from contrasting each stimulation condition against a control stimulation. Future research is warranted to confirm task as well as stimulation specific results in comparison to baseline conditions.

Additionally, it is worth mentioning that only the interaction effect observed within the sensorimotor putamen as well as the task and stimulation effects reported in the posterior hippocampus survived correction for multiple comparisons. Results observed in the DLPFC and the (entire) hippocampus did not survive such correction and therefore need to be interpreted cautiously.

## Conclusions

The results of the present study indicate that both motor sequence learning and prefrontal stimulation can modulate multivoxel response patterns in deep brain regions that are critical for the motor sequence learning process. They also suggest that stimulation influenced early motor memory consolidation processes during wakefulness as evidenced by a stimulation-induced modulation of the reinstatement of task pattern into rest immediately following motor learning.

## Data Availability

The ethical approval granted by the local ethics committee does not permit the publication of data online.

## Acknowledgements

This work was supported by the Belgian Research Foundation Flanders (FWO; G099516N) and internal funds from KU Leuven. GA also received support from FWO (G0D7918N, G0B1419N, 1524218N) and Excellence of Science (EOS, 30446199, MEMODYN, with SPS and DM). MAG, ND and MPV received salary support from these grants. MAG is funded by a predoctoral fellowship from FWO (1141320N). Financial support for author BRK was provided by the European Union’s Horizon 2020 research and innovation program under the Marie Skłodowska-Curie grant agreement (703490) and a postdoctoral fellowship from FWO (132635). EMR received salary support from the Air Force Office of Scientific Research (AFOSR, Virginia, USA; FA9550-16-1-0191). We wish to thank Kaat van Rooij and Michelle Roussard for assistance with data collection as well as Liesbeth Bruckers and Steffen Fieuws for assistance with statistical analyses.

## Author contributions

**Mareike Gann**: Conceptualization; Resources; Data curation; Software; Formal analysis; Investigation; Visualization; Writing - original draft; Writing - review and editing **Bradley King**: Conceptualization; Resources; Data curation; Software; Formal analysis; Validation; Writing - review and editing **Nina Dolfen**: Investigation; Methodology; Writing - review and editing **Menno Veldman**: Investigation; Writing - review and editing **Marco Davare**: Conceptualization; Methodology; Writing - review and editing **Stephan Swinnen**: Methodology; Writing - review and editing **Dante Mantini**: Resources; Software; Methodology; Writing - review and editing **Edwin Robertson**: Conceptualization; Writing - review and editing **Genevieve Albouy**: Conceptualization; Resources; Data curation; Software; Formal analysis; Supervision; Funding acquisition; Validation; Investigation; Writing - original draft; Project administration; Writing - review and editing

## Additional information

The authors have no conflict of interest to declare.

