## Supplemental Information for "Prefrontal stimulation prior to motor sequence learning alters multivoxel patterns in the striatum and the hippocampus"

#### Motor sequence learning and prefrontal repetitive TMS modulate multivoxel patterns in the DLPFC, hippocampus and putamen

Mareike A. Gann, Bradley R. King, Nina Dolfen, Menno P. Veldman, Marco Davare, Stephan P. Swinnen, Dante Mantini, Edwin M. Robertson, Geneviève Albouy

### Supplemental Tables

**Supplemental Table S1: Distribution of the different conditions per visit of all included conditions.**

| | Visit 1 | Visit 2 | Visit 3 | Visit 4 |
|---|---|---|---|---|
| **cSEQ** | 2 | 4 | 5 | 7 |
| **cRND** | 6 | 3 | 5 | 4 |
| **iSEQ** | 5 | 7 | 4 | 3 |
| **iRND** | 5 | 5 | 5 | 3 |

*The distribution of the different conditions per visit of all included conditions was misbalanced due to data exclusion (see main text for details) after data were acquired with a balanced procedure.*

**Supplemental Table S2: Individual TBS targets in MNI space.**

| Participant | x mm | y mm | z mm |
|---|---|---|---|
| P1 | -36 | 16 | 36 |
| P2 | -20 | 30 | 44 |
| P3 | -18 | 18 | 40 |
| P4 | -16 | 18 | 48 |
| P5 | -28 | 16 | 42 |
| P6 | -24 | 16 | 36 |
| P7 | -20 | 20 | 40 |
| P8 | -28 | 18 | 54 |
| P9 | -36 | 16 | 36 |
| P10 | -34 | 16 | 42 |
| P11 | -22 | 26 | 42 |
| P12 | -28 | 24 | 38 |
| P13 | -28 | 32 | 44 |
| P14 | -28 | 18 | 36 |
| P15 | -18 | 24 | 48 |
| P16 | -38 | 12 | 54 |
| P17 | -28 | 12 | 38 |
| P18 | -18 | 14 | 52 |
| P19 | -20 | 12 | 46 |

**Supplemental Table S3. Means and means adjusted for TBS visit effect.**

| ROI | SI | means | | | | means adjusted for TBS visit | | | |
|---|---|---|---|---|---|---|---|---|---|
| | | *cSEQ* | *cRND* | *iSEQ* | *iRND* | *cSEQ* | *cRND* | *iSEQ* | *iRND* |
| DLPFC | Early\|Late | .835 | .852 | .814 | .899 | .834 | .851 | .819 | .9 |
| DLPFC | RSpre\|RSpost | .621 | .654 | .64 | .631 | .629 | .647 | .642 | .627 |
| DLPFC | Task\|RSpost | .501 | .534 | .518 | .521 | .497 | .536 | .521 | .522 |
| Hippocampus | Early\|Late | .282 | .29 | .285 | .29 | .279 | .291 | .285 | .291 |
| Hippocampus | RSpre\|RSpost | .182 | .192 | .192 | .195 | .183 | .193 | .192 | .195 |
| Hippocampus | Task\|RSpost | .128 | .125 | .126 | .127 | .129 | .124 | .126 | .127 |
| Associative Putamen | Early\|Late | .486 | .497 | .483 | .473 | .484 | .5 | .481 | .474 |
| Associative Putamen | RSpre\|RSpost | .395 | .41 | .411 | .421 | .396 | .41 | .41 | .421 |
| Associative Putamen | Task\|RSpost | .286 | .275 | .288 | .285 | .284 | .276 | .288 | .286 |
| Sensorimotor Putamen | Early\|Late | .463 | .478 | .477 | .465 | .463 | .479 | .477 | .465 |
| Sensorimotor Putamen | RSpre\|RSpost | .430 | .443 | .436 | .443 | .412 | .44 | .437 | .442 |
| Sensorimotor Putamen | Task\|RSpost | .294 | .288 | .284 | .292 | .294 | .289 | .284 | .292 |
| Anterior Hippocampus | Early\|Late | .361 | .372 | .363 | .371 | .36 | .372 | .364 | .372 |
| Anterior Hippocampus | RSpre\|RSpost | .242 | .257 | .254 | .254 | .242 | .258 | .254 | 254 |
| Anterior Hippocampus | Task\|RSpost | .169 | .168 | .172 | .169 | .17 | .167 | .171 | .169 |
| Posterior Hippocampus | Early\|Late | .452 | .457 | .45 | .45 | .449 | .459 | .449 | .451 |
| Posterior Hippocampus | RSpre\|RSpost | .351 | .362 | .37 | .374 | .348 | .366 | .37 | .376 |
| Posterior Hippocampus | Task\|RSpost | .254 | .243 | .24 | .244 | .255 | .242 | .24 | .243 |

*Unadjusted means were derived from the linear mixed models not including the TBS visit as fixed effect. SI – similarity index, ROI – region of interest, TBS – theta-burst stimulation, DLPFC – dorsolateral prefrontal cortex, RS – resting-state, I – intermittent, c – continuous, SEQ – sequence, RND – random.*

**Supplemental Table S4. Task by stimulation effects on pattern similarity of the sub-territories of the hippocampus between RS pre- and post-stimulation/task controlling for the visit effect.**

| | dfs | F | *p* |
|---|---|---|---|
| **Anterior Hippocampus** | | | |
| *Task effect* | 1,18.930 | 2.926 | .104 |
| *Stimulation effect* | 1,17.090 | .895 | .357 |
| *Interaction* | 1,18.086 | 1.325 | .280 |
| **Posterior Hippocampus** | | | |
| *Task effect* | 1,18.879 | 6.239 | **.022*** |
| *Stimulation effect* | 1,11.878 | 7.620 | **.017*** |
| *Interaction* | 1,16.916 | 1.011 | .329 |

*Bold values indicate p<.05. False-discovery-rate (FDR) correction for multiple comparisons (2 ROIs) was applied with the Benjamini-Hochberg procedure (Benjamini and Hochberg, 1995). Asterisk (*) indicates significance at $p_{FDR}$<.05. dfs – degrees of freedom.*

**Supplemental Table S5. Task by stimulation effects on pattern similarity between task practice and RS post-stimulation/task controlling for the visit effect.**

| | dfs | F | *p* |
|---|---|---|---|
| **Anterior Hippocampus** | | | |
| *Task effect* | 1,19.064 | .460 | .506 |
| *Stimulation effect* | 1,17.256 | .200 | .660 |
| *Interaction* | 1,18.225 | .003 | .954 |
| **Posterior Hippocampus** | | | |
| *Task effect* | 1,15.996 | 1.772 | .202 |
| *Stimulation effect* | 1,15.074 | 10.336 | **.006*** |
| *Interaction* | 1,16.333 | 4.535 | **.049** |

*Bold values indicate p <.05. Asterisk (*) indicates significance at $p_{FDR}$<.05. dfs – degrees of freedom.*

#### Supplemental Figures

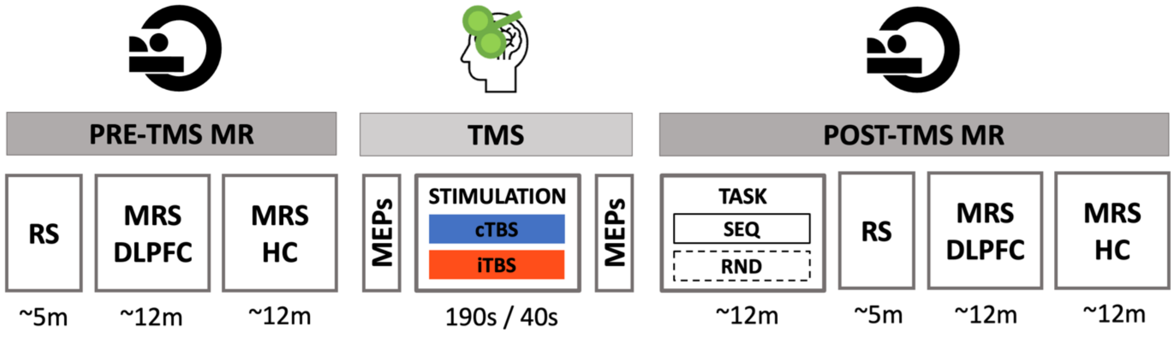


**Supplemental Figure S1. Full experimental design.** In each experimental session, participants first underwent pre-TMS whole-brain resting-state (RS) fMRI scans and magnetic resonance spectroscopy (MRS) scans of the dorsolateral prefrontal cortex (DLPFC) and the hippocampus (HC) that were followed by T1-neuronavigated intermittent or continuous theta-burst stimulation (iTBS or cTBS) applied to an individually-defined DLPFC target outside the scanner. Motor evoked potentials (MEPs) were measured pre- and post-TBS to probe corticospinal excitability. Immediately following the end of the TMS session, participants were placed in the MR scanner where they were trained on the motor task (sequential [SEQ] or random [RND] versions of the serial reaction time task) while BOLD images were acquired. After task completion, post-TBS/task RS and MRS data of the DLPFC and hippocampus were acquired. The order of the four experimental conditions in this within-subject design [cTBS/SEQ (cSEQ), cTBS/RND (cRND), iTBS/SEQ (iSEQ), iTBS/RND (iRND)] was counterbalanced across participants. Note that the data related to the MRS scans are not reported in the present manuscript. TMS: transcranial magnetic stimulation. [Figure adapted from (Gann et al., 2021)].

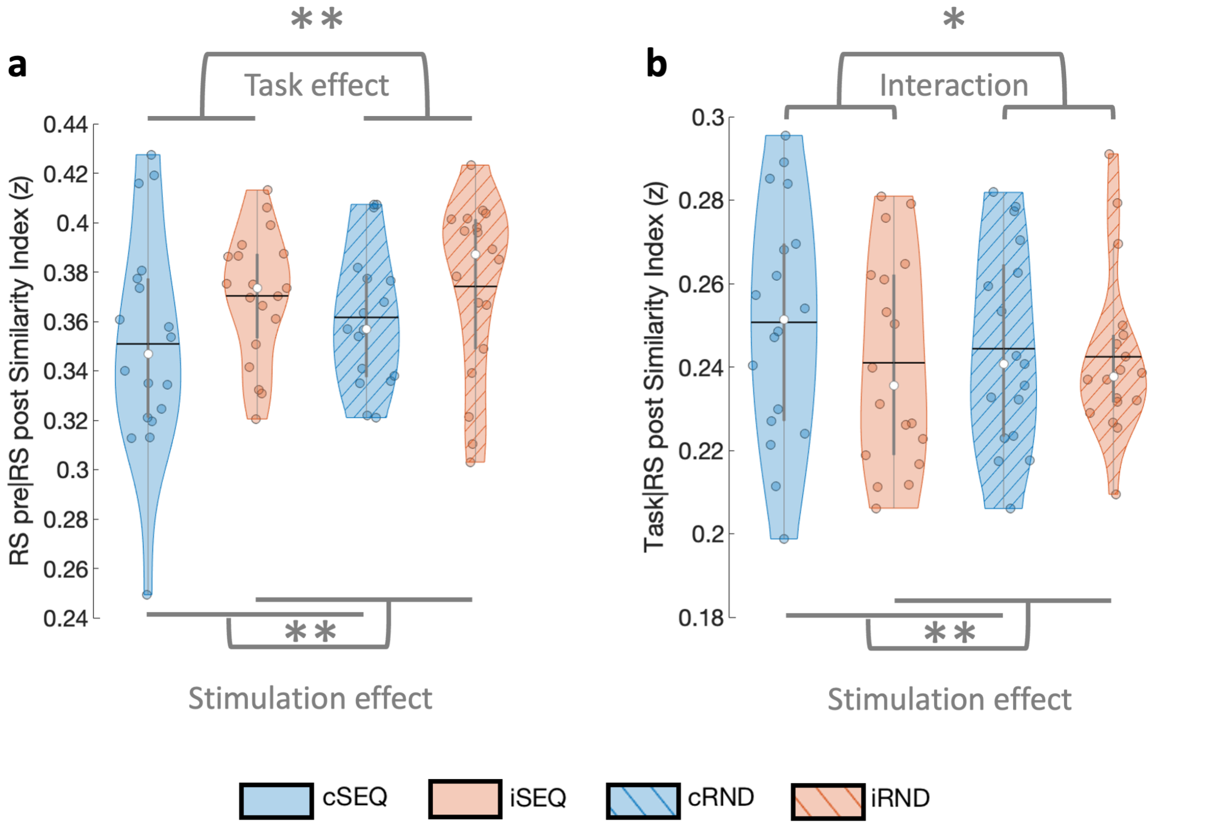


**Supplemental Figure S2. Posterior hippocampus patterns.** (a) Pattern similarity between pre and post RS was influenced by task and stimulation condition such that lower similarity was observed after sequence learning as compared to random practice as well as after cTBS as compared to iTBS. (b) Pattern similarity between task and RS post was influenced by stimulation condition such that higher similarity was observed after cTBS as compared to iTBS. The interaction effects were driven by higher similarity after cSEQ as compared to iSEQ. Colored circles represent individual data, jittered in arbitrary distances on the x-axis within the respective violin plot to increase perceptibility. Black horizontal lines represent means and white circles represent medians. The shape of the violin plots depicts the distribution of the data and grey vertical lines represent quartiles. Asterisk indicates significance at $p<.05$ (*) and at $p_{FDR}<.05$ (**). RS – resting-state, SEQ – sequence learning task version, RND – random task version, c – continuous, i – intermittent.
